# Experience reduces route selection on conspecifics by the collectively migrating white stork

**DOI:** 10.1101/2023.11.21.567993

**Authors:** Hester Brønnvik, Elham Nourani, Wolfgang Fiedler, Andrea Flack

## Abstract

Migration can be an energetically costly behavior with strong fitness consequences in terms of mortality and reproduction ^1–11^. Migrants should select migratory routes to minimize their costs, but both costs and benefits may change with experience ^12–14^. This raises the question of whether experience changes how individuals select their migratory routes. Here we investigate the effect of age on route selection criteria in a collectively migrating soaring bird, the white stork (*Ciconia ciconia*). We perform step selection analysis on a longitudinal data set tracking 158 white storks over up to nine years to quantify how they select their routes based on the social and atmospheric environments, and to examine how this selection changes with age. We find clear ontogenetic shifts in route selection criteria. Juveniles choose routes that have good atmospheric conditions and high conspecific densities. Yet, as they gain experience storks’ selection on the availability of social information reduces—after their fifth migration experienced birds also choose routes with low conspecific densities. Thus, our results suggest that as individuals age, they gradually replace information gleaned from other individuals with information gained from experience, allowing them to shift their migration timing and increasing the time scale at which they select their routes.

## Results

Adults often outperform juveniles in exploiting resources ^15–18^ and in saving energy ^19,20^. The necessary skills for efficiently finding and using resources may be obtained early in life ^21–23^ and for migrants these skills may be vital for selecting low-cost migratory routes. Yet, how experience affects migratory efficiency remains poorly understood ^12,13,24^. Knowledge regarding where to go or how to move efficiently can come from individual experience through trial and error ^25^, but when obtaining this information asocially becomes costly, individuals may switch to exploiting social information ^26,27^. Quantifying how the social environment affects migratory decisions has proven difficult because of the vast spatial scale of many migrations and the difficulty of quantifying the presence of conspecifics ^28–30^. Animals can learn their migratory routes if they travel in the same time and place as informed conspecifics ^31–35^. The age of an individual may also affect how it moves through its environment ^36–38^ and experienced individuals may be able to move more efficiently ^17,20^. But, it remains unclear how route choice is influenced by the social environment and whether this influence changes as individuals age.

To address this knowledge gap, we used white storks (*Ciconia ciconia*), collectively migrating, soaring birds, as a model. Flapping flight is energetically expensive for soaring birds ^39,40^, thus they cannot afford to migrate without riding uplifts and must choose routes that provide them ^41,42^. Inexperienced soaring birds may not be able to locate the invisible uplifts they need to plot the most energetically favorable routes. In this case, social information regarding when and where uplifts are available may be valuable ^43,44^. Social information can also help birds locate suitable wintering habitats, breeding sites, or movement corridors ^30,35,45^. However, although we know of pulses of migrants in the air during certain times of the seasons ^46,47^, their impact on migratory route selection at the individual level has not been studied before.

## The social and energy availability landscapes predict migratory route selection

To understand how white storks select their migratory routes, we analyzed the trajectories of 158 individuals tracked hourly over up to nine consecutive years of migration (Table S1). We conducted step-selection analysis with generalized linear mixed models ^48^ using proxies of thermal uplift and the probability of social information being available in a given place and time, along with the number of years of experience, as predictors of whether a location was used or unused.

To proxy uplift, we calculated the convective velocity scale ^49^, which estimates the vertical velocity of air due to heating by the sun at a resolution of 0.25 degrees hourly (see supplemental code). For the availability of social information, we estimated the hourly utilization distributions of 397 migrating white storks on a 30 km grid ^50^ (Video S1). Because an animal’s optimal decisions may differ between seasons ^44,51–53^, we built separate models for fall and spring. Both models had good predictive performance; the fall model fit with a root mean square error (RMSE, a comparison of observed to predicted values) of 0.138 and the spring model fit with an RMSE of 0.140.

Route selection by white storks was influenced positively by the density of migrating conspecifics (p *<* 0.001, Figure 2, Table S2) regardless of uplift or of how many migrations an individual had completed. We also found a significant interaction between age and conspecific density, indicating that experience affects selection on conspecific density (p *<* 0.001, Figure 2 Conspecifics X Migrations). In addition, storks selected routes to maximize uplift at all conspecific densities and ages (fall p *<* 0.001 and spring p = 0.002, Figure 2). Age did not significantly affect this selection on uplift (fall p = 0.351 and spring p = 0.058, Figure 2).

**Figure 1.**
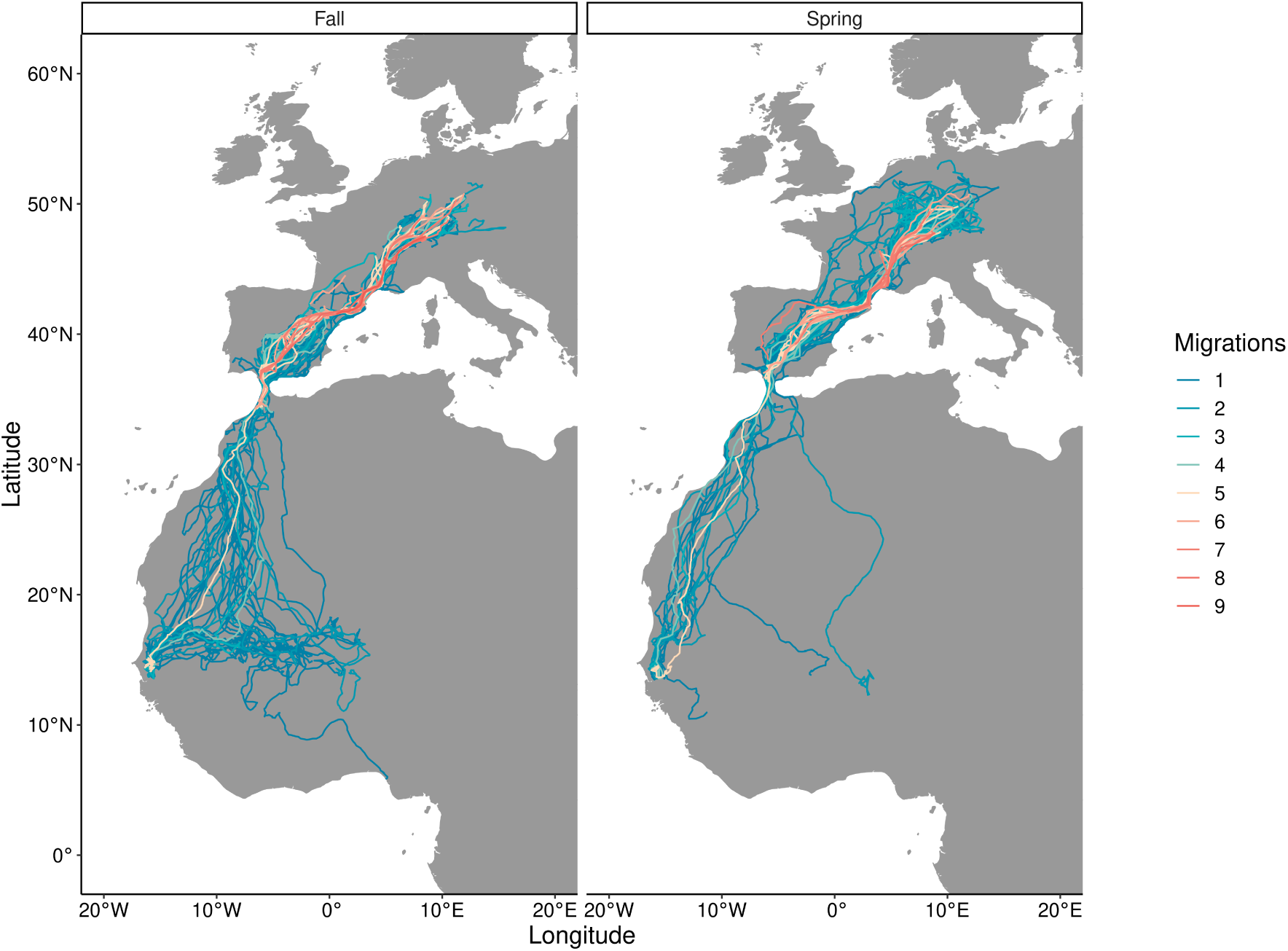
White storks travel diverse routes that vary between both seasons and ages. The migration routes of the 158 individuals included in our route-selection analysis. Colors indicate age, with each bird tagged in its nest and then tracked over up to nine years. See also Table S1.

**Figure 2.**
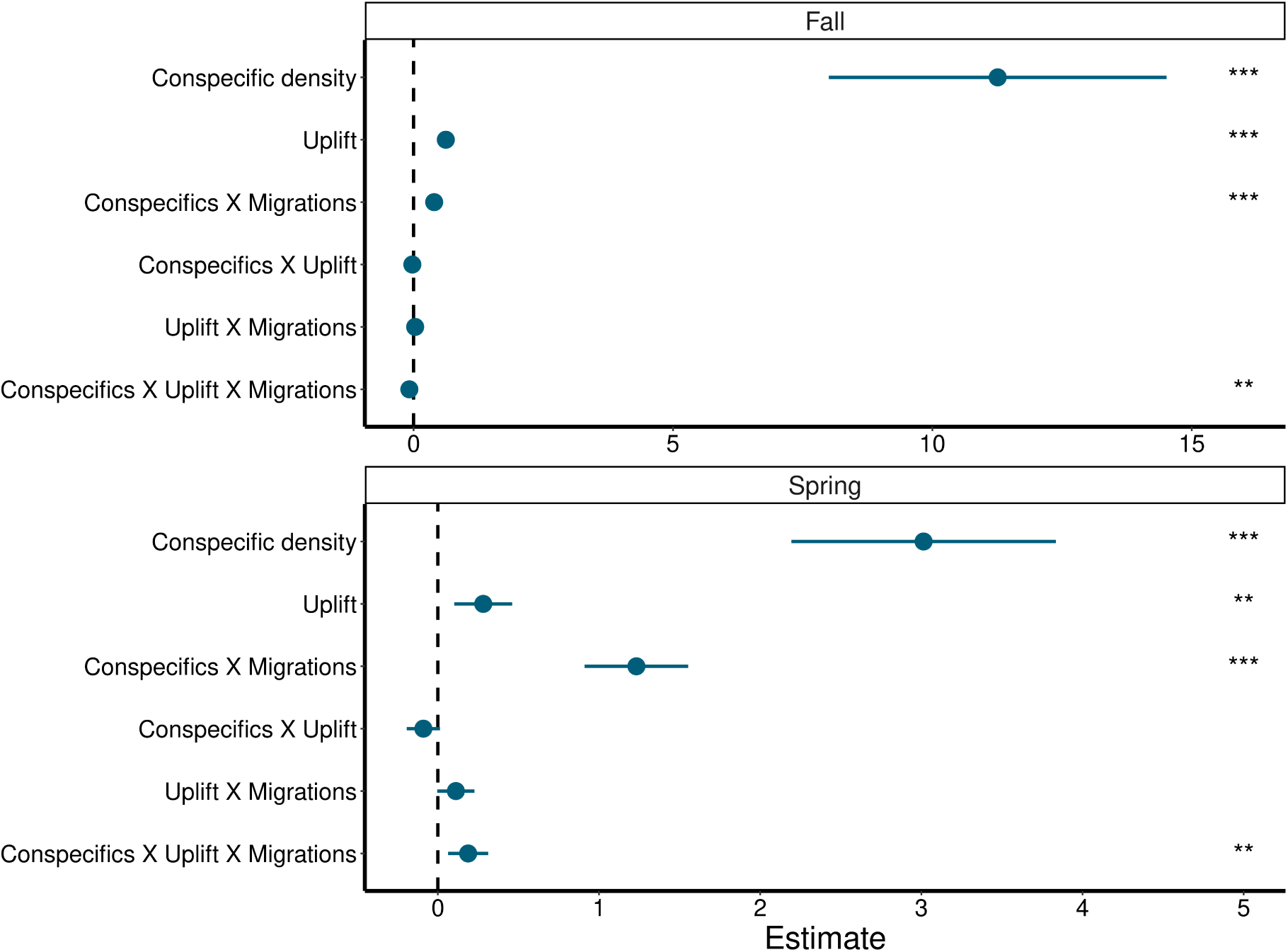
The interaction of conspecific density, uplift, and age predicts route selection. In both seasons, the availability of other storks, the availability of uplift, the interaction of conspecific density with age, and the interaction of all three variables are significant predictors of route selection. Posterior means (centered and scaled) and 95% credible intervals for the fixed effects in the GLMM for A) fall and B) spring migrations. Asterisks indicate the p-value ranges: p *<* 0.001 (***), p *<* 0.01 (**), p *<* 0.05 (*). See also Table S2.

### Age reduces the strength of migratory route selection on conspecific density

Next, to explore the significant interaction of conspecific density, uplift, and age, we used our fitted models to predict route selection across the spectrum of social and uplift conditions in each season.

Our models predicted that young birds selected for (selection probabilities *>* 0.50) high conspecific densities and against (selection probabilities *<* 0.50) low conspecific densities. But this preference shifted as birds aged, i.e. older birds were less likely to avoid low conspecific densities (Figure 3). This shift suggests that older birds were not as selective with regard to conspecific densities as younger birds were (Figure S2).

**Figure 3.**
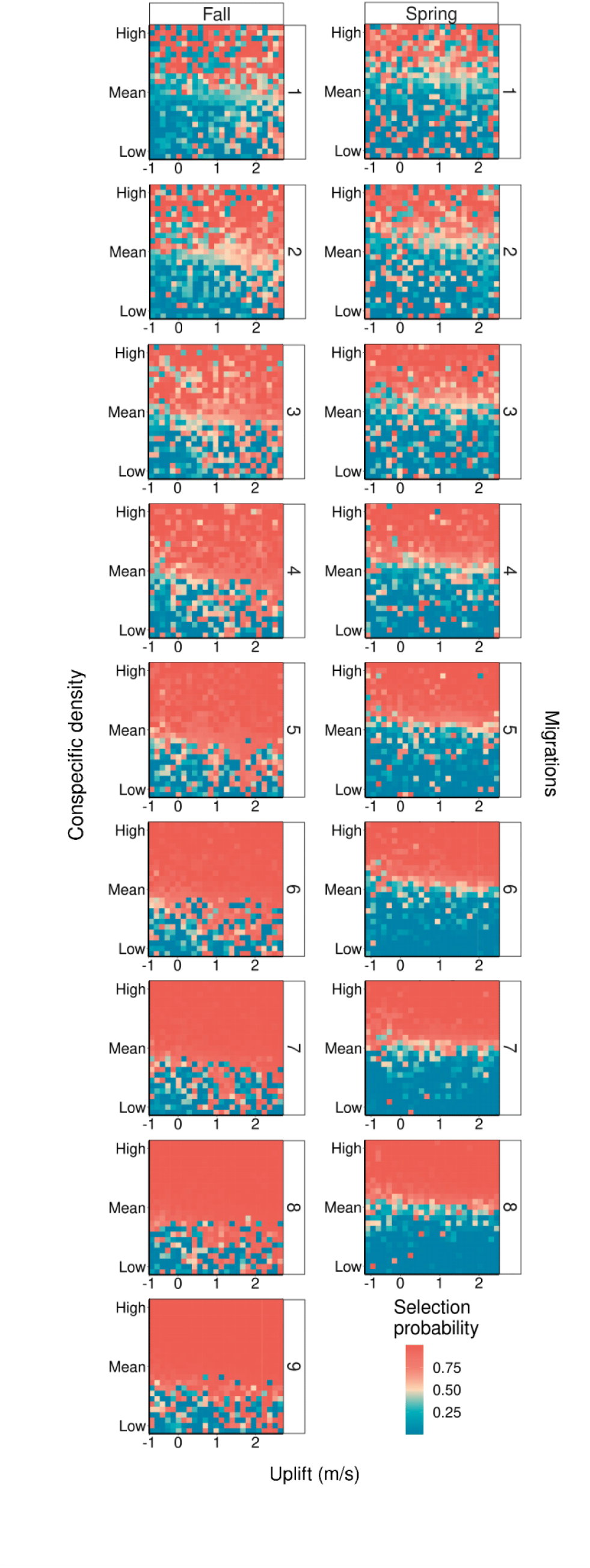
The strength of route selection depends on both age and season. Low selection probabilities (blue) indicate conditions that storks are predicted to select against (avoid), high selection probabilities (red) indicate conditions that the storks are predicted to use. Over consecutive migrations, white storks grew less selective on conspecific density at all uplift velocities (i.e., were less likely to select against low values). In fall, but not in spring, older birds became non-selective on conspecific density where strong uplift was available. See also Figure S2.

In fall, storks were predicted to have high selection probabilities (*>* 0.75) at low conspecific densities (below the mean) if strong uplift was available (*>* 1.5 m/s), thus in the fall storks will select for favorable atmospheric conditions even when doing so means experiencing lower conspecific densities. This pattern was not present in spring, indicating that in the spring storks will select for favorable atmospheric conditions, but only given favorable conspecific densities. This seasonal difference may be because in fall, saving time or reaching specific wintering grounds is less important than saving energy, or because uplift conditions deteriorate in spring (Figure S2) and increase the importance of conspecific density.

## Migration timing shapes the social and energy landscapes

As white storks age, they travel straighter, less exploratory routes ^13^ and shift their migration timing (Figure 4 A), which may mean that individuals of different ages do not experience the same environmental conditions. The observed strength of selection depends not just on the storks’ decisions but also on how variable their environmental conditions were. Thus, we may see a lack of selectivity due to true absence of selection by the individual and/or due to an absence of options provided by its environment. For this reason, we tested how much choice individuals of each age group had in which conditions to use.

**Figure 4.**
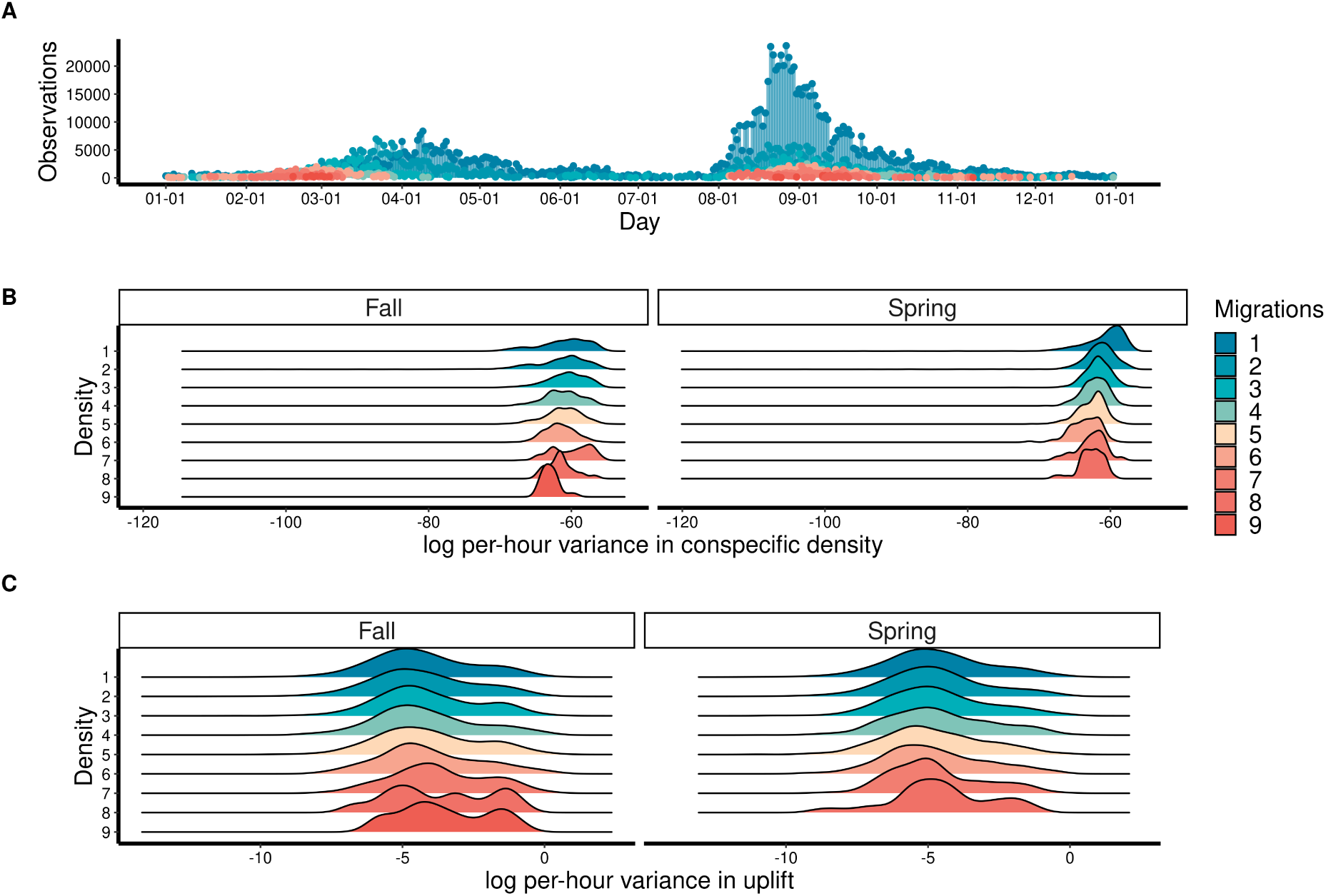
Different ages experience different environmental variability. A) Younger birds tended to migrate earlier in the fall and later in the spring than older birds did. B) We compared the variance per stratum in the data set (i.e. the variance in each set of one observed and 50 available locations in an hour) across ages using Kolmogorov-Smirnoff tests. The variance in available conspecific density reduced as storks aged. C) The variance in available uplift conditions increased as storks aged. See also Table S3.

We took the variance in the environment at each hour (each stratum in the step-selection model) and used Kolmogorov-Smirnoff tests to detect differences between the distribution of per-hour variance in each migration and the distribution of all other migrations’ per-hour variances pooled.

When comparing stratum level variance in conspecific density, i.e. the variance in the conditions available to an individual at each time point, we found it to be significantly lower for older birds (Figure 4 A & B, Table S3). As they aged, storks experienced decreasing variability. Thus, older birds had fewer choices and their reduced one-hourly selectivity might have been due to absence of options.

We found the opposite to be true of uplift. Older birds experienced increasing variability (Figure 4 C, Table S3). This variability could mean that older storks had to work harder to find rewarding patches, but also that they had an opportunity to be selective that the younger birds did not. Yet, we did not find a significant effect of age on selection of uplift (Figure 2).

As storks age they gain independence from flocking, as demonstrated by their reduced selection on conspecific density. The conspecific density available to adults was more uniformly distributed (Figure 4 B), indicating that they were less likely to congregate in large numbers. Thus, adults migrate at times when it may be more difficult to locate high conspecific densities. As storks age, they also improve in their ability to cope with sub-optimal uplift conditions— although all ages select on the availability of uplift, older birds have more heterogeneous uplift landscapes compared to the more predictable ones of younger birds. This pattern in uplift variability across age groups may be driven by differences in migration timing. Older storks tend to travel slightly later in the fall and earlier in the spring than younger ones do (Figure 4 A).

## Discussion

Here we investigated the effect of age on the use of access to social influences when selecting cost-efficient migration routes in a large soaring migrant. We quantified route selection by white storks at the one-hourly scale on the basis of conspecific density and uplift availability. We found that, as we predicted, storks select routes depending on uplift conditions but grew less selective on conspecific density with age as they repeated their migrations.

Conspecific density was an important predictor of route selection for all ages and we found a reduced strength of selection among older birds. The strength of selection depends on how different the environmental conditions are at the start and end of an hour-long step. The adults, which had lower selection strength, were experiencing more uniformly distributed conspecific density, thus their lower selectivity is a factor of fewer chances to be selective. The more uniform distribution of conspecific density available to older birds could be because: i) we have failed to capture the peaks and troughs of the social landscape due to dwindling sample size (Table S1, but see Methods), or ii) older storks are less likely to congregate in large numbers and really are adjusting their routes less based on where others are likely to be.

Older individuals may have a reduced dependence on locating flocks because their experience makes social information less valuable to them and allows them to make informed movement decisions independently. We observed a reduced selection on conspecifics at the scale of hourly locations. But, storks are also selecting when to migrate. In fall, experienced birds delay travel, ignoring the peak of the departures. In spring, they advance their return to the breeding area to rush ahead of their competitors, meaning that they may actually be avoiding traveling when there is the most social information and the most predictable uplift. Here we show that these choices at different temporal scales affect the availability of social and environmental support, revealing that storks may be selecting the conditions that they will have upon completing a migration rather than the instantaneous conditions they are experiencing en route. Thus, as animals age, they can make decisions at longer time scales informed not only by their perception of the current environment, but also by experience ^54^.

Although we found a strong effect of conspecific density on route selection, our estimates of migrating stork densities may be too narrow to capture all of the social hot-spots and their associated error will vary over space and time. The breeding white stork population is abundant in Europe and spreads from the southern borders of Spain up to southern Sweden, Denmark, and even the UK ^55^. We know that migration timing is different among these different sub-populations ^56^. These differences could have led us to underestimate the true availability of conspecifics because even if the birds in our study were unlikely to be traveling at a given time, there might have been birds from other populations that we could not monitor and that were in fact providing flocking opportunities to the tracked birds. Yet, it may be necessary to travel with others that are attempting to move in the same general direction. Thus, in spring it may be essential to perform return migrations with individuals from the same breeding population and our estimate of conspecific densities may capture these birds’ true experience and choices.

In general, young birds may need conspecifics not just for finding good flight conditions, but also to learn routes ^34,35,57^, to find breeding colonies ^58,59^, or to find suitable wintering grounds ^35^. Although here we only examined flight choices, these flights lead to suitable habitats. Thus, it is likely that these group flights lead to social learning that allows young birds to form spatial memories across scales that can be retrieved in the following years. These experiences may allow older, more experienced individuals to be less sensitive to social guidance but more sensitive to arrival timing, which may determine food availability in the winter and competition for nests in the spring. Effectively, this pattern means that older birds are selecting at longer time-scales, producing the observed reduction in selectivity at the one-hourly scale.

Route selection by migrants is influenced by many factors. For example, animals that rest and refuel during migration may need to consider available sleeping and foraging sites ^60^. Depredation during migration ^61^, disturbance ^62^, bad weather ^63,64^, and health ^65,66^ can also affect migratory decisions and may shape routes. When migrating repeatedly, spatial memory most likely also plays a significant role in migratory route decisions, which could be quantified using a previous year’s range as a predictor^67^. Here we show that one additional driver of the decisions of this social migrant was the social landscape and that its effect was altered by age.

In the trade-off that migration poses, older individuals can profit from past experience. We found that with age, white storks shifted their decisions away from the scale of an hour, most likely driven by the need to shift their travel times and arrival dates at the scale of weeks. Individuals of long-lived migratory species may travel to and from breeding grounds dozens of times. This repetition affords them the opportunity to learn on early migrations and use experience to improve later ones ^21^. Animals that travel in groups also have access to social information that can help them to negotiate their migrations and that may be of particular importance for inexperienced individuals ^35^. We show that white storks select their migratory routes to increase their overlap with areas that have a high probability of hosting other storks at that time. Selection reduces as they age and migrate when there are fewer other migrants. This indicates that experienced storks do not need to seek out potential flocks. We expect that individuals of other species that migrate in groups also need the benefits of social groups less as they gain experience in saving and gaining energy, avoiding danger, and navigating to their goals.

## Methods

### RESOURCE AVAILABILITY

#### Lead contact

Further information and requests for resources should be directed to and will be fulfilled by the lead contact, Hester Brønnvik.

#### Materials availability

This study did not generate new unique reagents.

#### Data and code availability

- All raw data necessary to replicate the findings of this study are available on Move-bank.org. Files containing only the processed data used in this study, including environmental covariates, will be made available upon acceptance of the manuscript.
- R scripts used for step selection analysis are available on *github* at: https://github.com/hesterbronnvik/stork_route_selection_public.git.
- Any additional information required to reanalyze the data reported in this paper is available from the lead contact upon request.

### EXPERIMENTAL MODEL AND SUBJECT DETAILS

White storks are long-distance, seasonal migrants. White storks breed on the Iberian Peninsula, and from central and eastern Europe to northwest Africa and western Asia ^56^, they then overwinter in Iberia and sub-Saharan Africa. White storks are soaring-gliding birds that rely on thermal uplift to offset their cost of transport ^56,63,68^. Migrating storks travel collectively in flocks ^43^ that may consist of hundreds of individuals ^63,69^. This could provide them with social information regarding routes ^40,43^ and the migration routes of experienced flock partners influence the routes of inexperienced individuals ^45^.

Flocks of migrating storks will consist largely of young birds as they are more represented in the population due to high early-life mortality rates. More specifically, a white stork breeding pair has approximately 3 chicks per year. The average life expectancy of a breeding bird is 8-10 years. With approximately 1000 breeding pairs in the German state of Baden-Württemberg (e.g. year 2016), ranging from age 3-30, the distribution of breeding birds is also biased towards younger birds. These breeding pairs produce around 3000 juveniles in a year. In addition, there is a substantial number of floaters (non-breeding birds, age 2-4) in the population, estimated to be around 1200 individuals per year (in our study area). After one year only 30% of the juveniles are still alive ^70^, and even two and three-year-old birds still have mortality rates of around 25%.

Permits for tagging and tracking the white storks were issued by the authorities of the Federal States (G-13/28 and G-268 15/47 by Regierungspräsidium Freiburg, 54-2532.1-14/14 by Regierung von Mittelfranken, MPI269 O-1/14 by Regierungspräsidium Tübingen, G15-20-032 by LUA Rheinland-Pfalz, ROB-55.2Vet-270 2532.Vet_02-17-95 by Regierung von Oberbayern).

### METHOD DETAILS

#### Movement data

We used existing data from several studies tracking white storks. In southwestern Germany, 464 juvenile white storks were tagged on the nest with solar GSM-GPS-ACC loggers between 2013 and 2022 (e-obs GmbH, Table S4, for details see ^13,56,70^).

We only analyzed complete, long distance migrations along the western flyway by birds from the southwest German population. We defined migration using simple speed thresholds. The start and end of a migration were the first and last days with a total displacement of at least 40 kilometers within a week of moving more than 70 kilometers in a day. This double threshold accounts for the fact that storks may move slowly as they depart and arrive. We defined complete, long-distance migrations as those that ended at least 30 days before the bird died and that displaced the bird at least 3 degrees of latitude. Using these definitions resulted in 502 routes taken by 158 birds after filtering (Table S1). Routes marked as complete may include those for which birds died on a stopover site because whether the bird intended to (or would have decided to) continue migration after 30 days cannot be known.

#### Step selection functions

Step selection analysis views movement as a series of discrete steps between locations ^71^. Each set of two locations is an observed step beginning and ending at observed locations. A set of alternative steps is generated beginning at the same observed location, but ending at alternative locations. These sets of one observed step and its alternatives are strata in the data set.

We down-sampled the birds’ locations to once per-hour, then we generated 50 alternative locations over land for each observed one. Our step lengths were randomly drawn from a gamma distribution that was fitted to all of the distances traveled during observed one-hour (*±* 15 minutes) flight segments, and our turning angles were randomly drawn from a von Mises distribution fitted to the observed turning angles (see Figure S1 and supplemental code. We annotated each location with the environmental variables associated with that hour.

#### Environmental variables

We used two environmental covariates in our analysis: thermal uplift velocity, and the density of migrating conspecifics (a proxy for availability of social information en route). We also investigated the effect of wind support, but did not include it in our final models as we we found no evidence supporting its importance (see supplementary code).

##### Availability of thermal uplift

To estimate the availability of thermal uplift on the landscape, we retrieved data from the European Centre for Medium-Range Weather Forecasts (ECMWF) ERA-5 Global Atmospheric Re-analysis (0.25 degree and 1 hourly resolution). We estimated uplift velocity as:

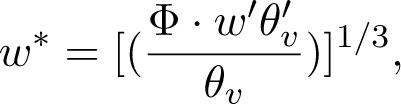

where Φ is the geopotential at the highest pressure level below the boundary layer height (m^2^ s*^−^*^2^), w’*θ_v_*’ is the kinematic heat flux (K m s*^−^*^1^), and *θ_v_* is the virtual potential temperature at the highest pressure level below the boundary layer height (K) ^41,49,72,73^ (for full calculations see supplementary code).

##### Availability of social information

To proxy the potential to find flock mates on the landscape, we estimated conspecific density as the utilization distributions of migrating storks. The distributions did not take into account the year, natal population, experience level, or whether the routes were complete. We estimated these utilization distributions from recorded locations using auto-correlated kernel density estimates (AKDEs) ^74,75^ (in the “ctmm” package ^50^). To make population-level distributions, we pooled the data from all storks that migrated from the southwest German population along with an additional 50 birds tagged in Spain, 79 in eastern Germany, and seven in France (397 migrated out of the 600, Table S4). We aligned the migratory locations (all locations between the start and end of migration, including non-flight segments) to the day, month, and hour, generating 3028 unique groups each with up to three observations from each individual that transmitted in that hour (for example, 12-21 12:00:00 to 12:59:00 UTC). To keep these distributions consistent and comparable, we built each AKDE on a 30 km x 30 km grid extending from 0*^◦^* to 60*^◦^* latitude and -20*^◦^* to 20*^◦^* longitude (using the Mollweide equal-area projection). However, the confidence did vary between distributions because the sample sizes were not the same in each hour (between 26 and 1428). Sample sizes differed particularly because there are unequal numbers of birds migrating at different parts of the year and because mortality meant our sample sizes were higher in the fall (Figure 4 A, Table S1). We generated one AKDE for each hour of the migrations of the 158 southwest German birds (see Video S1). Finally, we annotated each location with the average probability density of the grid cell where it fell (see supplementary code).

Our estimate of conspecific density is biased towards younger individuals. However, we are also confident that this is a realistic representation of the white stork’s proportions during migration ^69,76–78^. As detailed above (see Subject Details), there are large numbers of juveniles and high mortality rates. Thus, our estimated density is just a sample of the entire population but its temporal development (i.e. decrease of older birds in the sample) is also what happens in the population throughout time. In addition, we have included birds from more northern and southern populations (juveniles & older birds) to account for different departure times of other sub-populations.

### QUANTIFICATION AND STATISTICAL ANALYSIS

We performed all statistical analyses in *R* 4.3.1^79^, with criteria for significance set at p < 0.05. Sample size and the other relevant statistical details are provided in the supplemental tables.

#### Step selection functions

We estimated step-selection functions using generalized linear mixed models (using the *glmmTMB* package) ^48,80^. To ask how the roles of group availability and uplift in route selection change over time, we used conspecific density, uplift, a bird’s experience level (1-9 migrations), and the interactions of the three as predictors of use case (used location or available location). We standardized the predictor variables across the whole dataset by calculating z-scores. We included individual as a random effect on the slopes. Finally, we assessed model fit using root mean square error (RMSE) ^81^, a comparison of the observed value to values predicted by the model. Small values of RMSE indicate good predictive ability.

To know how the interaction of the predictors affected route selection, we used our fitted models to predict selection across the spectrum of potential environments. We created dummy data with 25 values evenly distributed across the middle 97.5% of the true uplift data (to exclude outliers) and 25 values evenly distributed across the middle 97.5% of the true conspecific density data separately for each season. Then, we used the models fitted to the actual data set to predict the probabilities of selection for each set of conditions in the dummy data. This provided us with predictions of how likely each age group was to use every combination (n = 625) of atmospheric and social conditions in each season (see supplementary code).

#### Environmental variability

The strength of selection is the difference between conditions along the observed and available steps ^82,83^. Thus, the strength of selection that we observe depends on the heterogeneity of the available conditions at each step. We explored whether the conditions that each age group experienced were equally heterogeneous by comparing the distributions of variance in each variable, each season, and each migration. First, we took the variance in the environment at each step (one observed and 50 available locations, a stratum in the model). Then, we used Kolmogorov-Smirnoff tests to detect differences between the distribution of per-stratum variance in each migration and the distribution of all other migrations’ per-stratum variances pooled.

**Legend for Video S1. The estimated conspecific density of migrating white storks (*Ciconia ciconia*), Related to STAR Methods.** We pooled the data from 397 white storks tagged in Germany (eastern n = 15, western n = 345), France (n = 5), and Spain (n = 32) between 2013 and 2022. If an individual was migrating in a given hour, we included its data in an autocorrelated kernel density estimate (AKDE). Each AKDE yielded a utilization distribution from which we took the probability density function as an estimate of where and when white storks (and thus the social information they generate) are likely to be available to conspecifics.

## Acknowledgments

We thank the members of the Collective Migration and Animal-environment Interactions research groups at the Max Planck Institute of Animal Behavior for providing many constructive comments. Kamran Safi gave valuable advice on the theoretical framework. We also thank Martin Wikelski for his support, discussions, and feedback. We sincerely thank Christen Fleming and Björn Reineking for early input regarding analysis, Gil Bohrer for help with uplift estimates, and Klaus Reuter and Sarah Davidson for technical support. We would also like to thank Michael Kaatz, Julio Blas, and everyone who helped with the extensive tagging efforts in Germany, France, and Spain. Finally, we thank Will Oestreich and two anonymous reviewers for many helpful questions and comments. HB was supported by the German Research Foundation (DFG, Emmy Noether Fellowship 463925853 to AF) and the International Max Planck Research School for Quantitative Behaviour, Ecology and Evolution (IMPRS-QBEE). EN was supported by the PRIME programme of the German Academic Exchange Service (DAAD) with funds from the German Federal Ministry of Education and Research (BMBF). AF was supported by the German Research Foundation (DFG, Emmy Noether Fellowship 463925853), the Hans und Helga Maus-Stiftung, and the James Heineman research award of the Minerva Stiftung. This work was supported by the Max Planck Society.

## Author contributions

H.B.: formal analysis, investigation, methodology, visualization and writing—original draft; E.N.: conceptualization, methodology, validation and writing—review and editing; W.F. resources, funding acquisition, and writing—review and editing; A.F.: methodology, validation, resources, funding acquisition, supervision, and writing—review and editing.

All authors gave final approval for publication and agreed to be held accountable for the work performed therein.

