## Supplemental Figures and Tables for "Experience reduces route selection on conspecifics by the collectively migrating white stork"

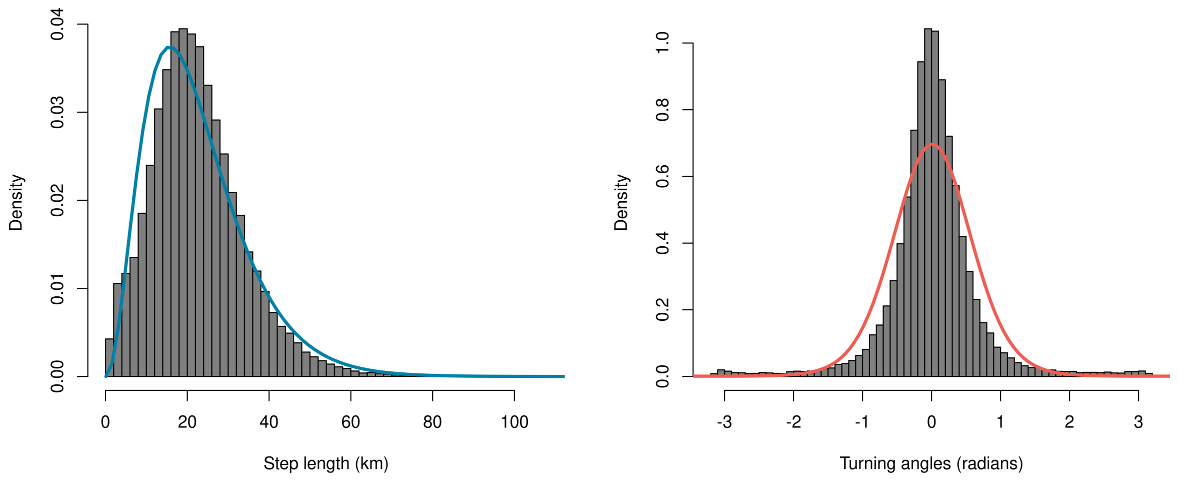


**Figure S 1 . Available locations are generated using the observed step lengths and turning angles and observed locations to define availability, Related to the STAR Methods.** The distributions of observed (histograms) and alternative (densities in color) step lengths (left) and turning angles (right). We randomly drew step lengths and turning angles from the alternative distributions fitted to the observed values and used them to generate steps available to the bird given its location and abilities.

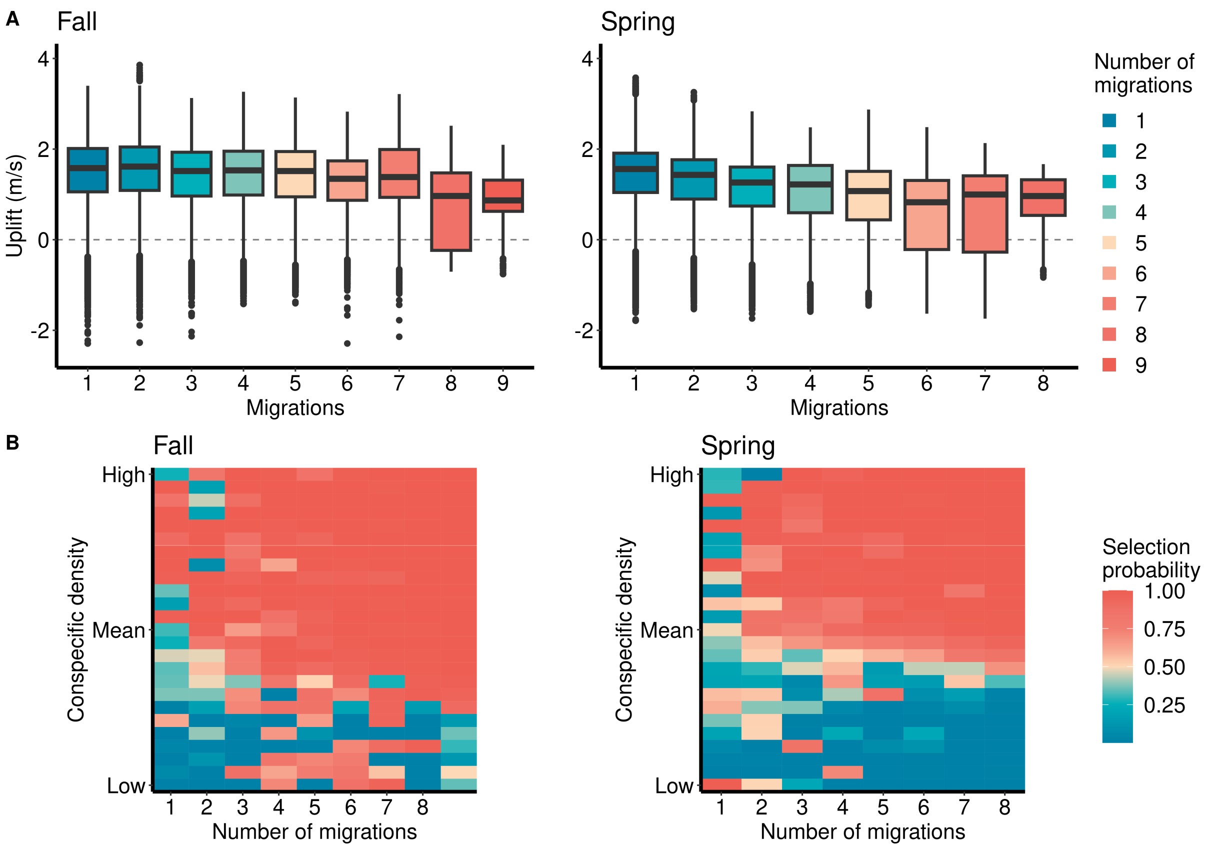


**Figure S 2 . Older birds do not select against low conspecific densities as strongly as young ones do, Related to Figure 3.** A) The distributions of available uplift (convective velocity scale) for birds of different ages in each season. B) The predictions from generalized linear mixed models holding uplift at its seasonal mean and exploring the significant interaction of number of completed migrations with conspecific density. Low selection probabilities (blue) indicate conditions that storks are predicted to select against (avoid), high selection probabilities (red) indicate conditions that the storks are predicted to use. Over consecutive migrations, white storks were less likely to select against low conspecific densities.

| Migration | Fall | Spring |
|---|---|---|
| 1 | 157 | 76 |
| 2 | 50 | 53 |
| 3 | 33 | 28 |
| 4 | 18 | 16 |
| 5 | 17 | 14 |
| 6 | 11 | 11 |
| 7 | 8 | 6 |
| 8 | 2 | 1 |
| 9 | 1 | 0 |

**Table S 1 . The number of tracks used for step-selection analysis, Related to Figure 1 and the STAR Methods.** After filtering for migrants along the western flyway that survived at least one migration, we had data from 158 individuals. One of those did not contribute its first fall migration, but did contribute a first spring and second fall, accounting for the difference between number of individuals and number of tracks.

| Variable | Estimate | Upper | Lower | p-value |
|---|---|---|---|---|
| **Fall** | | | | |
| Conspecific density | 11.3 | 14.5 | 8.00 | 0.000 |
| Uplift | 0.621 | 0.708 | 0.534 | 0.000 |
| Conspecifics X Migrations | 0.396 | 0.563 | 0.229 | 0.000 |
| Conspecifics X Uplift | -0.025 | 0.025 | -0.074 | 0.325 |
| Uplift X Migrations | 0.030 | 0.093 | -0.033 | 0.351 |
| Conspecifics X Uplift X Migrations | -0.082 | -0.031 | -0.133 | 0.002 |
| **Spring** | | | | |
| Conspecific density | 3.01 | 3.83 | 2.19 | 0.000 |
| Uplift | 0.282 | 0.461 | 0.102 | 0.002 |
| Conspecifics X Migrations | 1.23 | 1.55 | 0.910 | 0.000 |
| Conspecifics X Uplift | -0.090 | 0.013 | -0.193 | 0.088 |
| Uplift X Migrations | 0.112 | 0.227 | -0.004 | 0.058 |
| Conspecifics X Uplift X Migrations | 0.188 | 0.312 | 0.064 | 0.003 |

**Table S 2 . The model summary for the generalized linear mixed models predicting route selection in each season, Related to Figure 2.** We used the interaction of conspecific density, convective velocity scale, and the number of migrations an individual had completed as predictors of whether a location was used or unused in GLMMs.

| | Uplift | | | Conspecific density | | |
|---|---|---|---|---|---|---|
| Migration | Test statistic | p-value | sig. | Test statistic | p-value | sig. |
| **Fall** | | | | | | |
| 1 | 0.028 | 0.0002 | yes | 0.075 | 0 | yes |
| 2 | 0.020 | 0.1096 | no | 0.068 | 0 | yes |
| 3 | 0.046 | 0.0006 | yes | 0.085 | 0 | yes |
| 4 | 0.027 | 0.4639 | no | 0.095 | 0 | yes |
| 5 | 0.045 | 0.0414 | yes | 0.115 | 0 | yes |
| 6 | 0.063 | 0.0226 | yes | 0.199 | 0 | yes |
| 7 | 0.163 | 0 | yes | 0.160 | 0 | yes |
| 8 | 0.196 | 0.101 | no | 0.270 | 0.0069 | yes |
| 9 | 0.230 | 0.036 | yes | 0.538 | 0 | yes |
| **Spring** | | | | | | |
| 1 | 0.035 | 0.0005 | yes | 0.323 | 0 | yes |
| 2 | 0.035 | 0.0013 | yes | 0.122 | 0 | yes |
| 3 | 0.015 | 0.8866 | no | 0.168 | 0 | yes |
| 4 | 0.044 | 0.1403 | no | 0.241 | 0 | yes |
| 5 | 0.049 | 0.1070 | no | 0.321 | 0 | yes |
| 6 | 0.059 | 0.1670 | no | 0.410 | 0 | yes |
| 7 | 0.080 | 0.2582 | no | 0.302 | 0 | yes |
| 8 | 0.123 | 0.5686 | no | 0.349 | 0.0001 | yes |

**Table S 3 . None of the distributions of variances of conspecific density matched the distributions of the others, Related to Figure 4.** The distributions of variances in both available conspecific density and available uplift increasingly differ from the others in the fall. Asymptotic two-sample Kolmogorov-Smirnov tests comparing the distribution of each migration's per-stratum variance in environment to the distribution of all the other migrations pooled.

| Study | Individuals | Tracks | Analyzed | DOI |
|---|---|---|---|---|
| 24442409 | 67 | 45 | 18 | N/A |
| 212096177 | 18 | 14 | 9 | https://www.doi.org/10.5441/001/1.c42j3js7 |
| 76367850 | 108 | 78 | 37 | https://www.doi.org/10.5441/001/1.4192t2j4 |
| 21231406 | 92 | 78 | 32 | https://www.doi.org/10.5441/001/1.ck04mn78 |
| 1176017658 | 153 | 113 | 54 | https://www.doi.org/10.5441/001/1.277 |
| 173641633 | 26 | 17 | 8 | https://www.doi.org/10.5441/001/1.71r7pp6q |
| 1562253659 | 7 | 5 | 0 | N/A |
| 9648615 | 42 | 25 | 0 | N/A |
| 2204484313 | 38 | 11 | 0 | N/A |
| 10449318 | 41 | 4 | 0 | N/A |
| 2106043019 | 8 | 7 | 0 | N/A |

**Table S 4 . The sources of the tracking data, Related to the STAR Methods.** The identifier of each Movebank study, number of individuals tracked in that study, number of migratory tracks in the study, and how many tracks we used for route selection analyses.
